# Taking time to compose thoughts with prefrontal schemata

**DOI:** 10.1101/2023.07.25.550523

**Authors:** Kwang Il Ryom, Anindita Basu, Debora Stendardi, Elisa Ciaramelli, Alessandro Treves

## Abstract

Under what conditions can prefrontal cortex direct the composition of brain states, to generate coherent streams of thoughts? Using a simplified Potts model of cortical dynamics, crudely differentiated into two halves, we show that once activity levels are regulated, so as to disambiguate a single temporal sequence, whether the contents of the sequence are mainly determined by the frontal or by the posterior half, or by neither, depends on statistical parameters that describe its microcircuits. The frontal cortex tends to lead if it has more local attractors, longer-lasting and stronger ones, in order of increasing importance. Its guidance is particularly effective to the extent that posterior cortices do not tend to transition from state to state on their own. The result may be related to prefrontal cortex enforcing its temporally-oriented schemata driving coherent sequences of brain states, unlike the atemporal “context” contributed by the hippocampus. Modelling a mild prefrontal (vs. posterior) lesion offers an account of mind-wandering and event construction deficits observed in prefrontal patients.

## 1 Constructive associative memories

Recent explorations of the mechanisms underlying *creative* forms of human cognition [1, 2], ranging from musical improvisation [3] through visual creativity [4] up to poetry [5], or mere mind wandering [6], have again questioned the validity of reducing the cortex to a machine operating a complex transformation of the input it currently receives. On the one hand, sophisticated and massive artificial intelligence systems like ChatGPT or midJourney, with their impressive performance, have adhered to the standard operational paradigm of producing a response to a query. On the other, a simple observation of cortical circuitry, with its extensive recurrence and quantitatively limited external inputs, have long ago led to the proposal that the cortex is (largely) a machine talking to itself [7]. Likewise, when confronted with an artistic or literary creation we sometimes ask: what was the query? Was there a query?

If it is the cortex itself that takes the initiative, so to speak, is it the *entire* cortex? Understanding the mechanisms of cortico-cortical dialogue that generate spontaneous behaviour cannot eschew their statistical character, that of a system with very many imprecisely interacting elements. Valentino Braitenberg suggested a framework for such a statistical analysis, which to a first approximation considers the cortex as a homogeneous structure, not differentiated among its areas (nor, other than quantitatively, among mammalian species) [8]: the only distinction is between long-range connections and local ones – those which reach in the immediate surround of the projecting neuron and do not travel through the white matter. Importantly, by asking whether there is any computational principle other than just associative memory operating at both long-range and local synapses [7], Braitenberg pushes the age-old debate of whether cortical activity is more like a classic orchestra led by a conductor or more like a jazz jam session, beyond the limits of abstract information-processing models. In traditional box-and-arrows models of that kind, a box, whether it represents a specific part of the brain or not, can operate any *arbitrary* transformation of its input, which makes it difficult to relate it to physiological measures, and tends to leave the debate ill-defined. If at the core one is dealing solely with associative memory, instead, the issue can be approached with well-defined formal models, generating statistical insights that can be later augmented with cognitive qualifications.

Given the canonical cortical circuit [9] as a basic wiring plan for the generic cortical plaquette, or patch, getting at the gist of how it contributes to the exchanges mediated by long-range cortico-cortical connectivity among different patches requires considering the fundamental aspects that vary, at least quantitatively, among the areas. A number of reviews [10, 11] have pointed out that several prominent features align their gradients of variation, across mammals and in particular in the human brain, along a *natural cortical axis*, roughly from the back to the front of the cortex. Actual observations and measurements may be incomplete or even at variance with such a sweeping generalization, but here we take it as a convenient starting point. Anatomical measures point at more spines on the basal dendrites of pyramidal cells, indicating more local synaptic contacts in temporal and especially frontal, compared to occipital cortex [12]. This may support a capacity for more and/or stronger local attractor states. More linear and prompt responses to afferent inputs in posterior cortices, e.g. visual ones [13, 14], also suggest reduced local feedback relative to more anterior areas.

The rapidity of the population response to an incoming input has been related to the notion of an intrinsic *timescale* that might characterize each cortical area, and that may produce highly non-trivial effects, for example when inhibiting a particular area with TMS [15]. The timescales measured with similar methods have been shown to differ considerably, even within individual areas [16], and to define distinct cortical *hierarchies*, when extracted in different behavioural states, e.g. in response to visual white noise stimuli [17] or during free foraging [18]. Thus it remains unclear whether the ambition to define a unique hierarchy of timescales can really be pursued [19], and whether they can be related to patterns of cortical lamination [20] and to biophysical parameters, including the *I*_*h*_ current and others underlying firing rates and firing frequency adaptation [21]. Still, in broad terms multiple timescale hierarchies do roughly align with the natural axis, from faster in the back to slower in the front of the brain, and ignoring a factor of, say, four [19] would appear to grossly overlook a basic principle of cortical organization. Here, we ask what are the implications of major differences in *cortical parameters* for how basic associative memory mechanisms may express cortically-initiated activity. We focus on a simple differentiation between a posterior and a frontal half of the cortex, and neglect finer distinctions, e.g., rostrocaudal hierarchies within prefrontal cortex [22, 23] or the undoubtedly major differences within posterior cortices.

## 2 A simply differentiated Potts model

The mathematically defined model we use is based on the abstraction of a network of 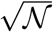 patches of cortex (where *𝒩* are all its pyramidal cells), interacting through long-range, associatively modified synapses, an abstraction close to that informing *connectome* research [24]. Each patch would be a densely interconnected network of 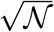 pyramidal cells interacting through local synapses, also associatively modifiable according to some form of Hebbian plasticity. Such a local cortical network may operate as an autoassociative memory once it has acquired through learning a number *S* of attractor states. In the simplified Potts formulation adopted here, the local network realized in each patch is replaced by a Potts unit with *S* states, and the analysis can focus on the network of long-range effective interactions between Potts units, which are no more mediated by simple synaptic connections, rather the connections are mathematically expressed as tensors [25].

We refer to previous studies [26] and to Appendix A for a description of the standard model, including the roles of the number *S* of local attractor states, of the feedback coefficient *w*, indicating how deep or strong they are, and of the adaptation time constant *τ*_2_, parametrizing the time it takes for the local network to be eased out of its current attractor.

A network of Potts units can express spontaneous behaviour when it *latches*, i.e., it hops from a quasi-stationary pattern of activity to the next, in the absence of external input – of a query [27]. Latching dynamics are a form of iterated associative memory retrieval; each extended activity pattern acts briefly as a global cortical attractor and, when destabilized by the rising thresholds which model firing rate adaptation, serves as a cue for the retrieval of the next pattern. Studies with brain-lesioned patients indicate, however, that there is structure in such spontaneous behaviour. In studies of mind-wandering, for example, patients with lesions to ventromedial prefrontal cortex (vmPFC) show reduced mind-wandering, and their spontaneous thoughts tend to be restricted, focused on the present and on the self, suggestive of a limited ability to project coherently into the future [28].

We then take our standard, homogeneous Potts network, differentiate it in two halves, and ask whether a structure of this type may reflect a basic differentiation between frontal and posterior cortices in the number or in the strength of their local attractor states, or in the time scale over which they operate, as expressed in differences, in the model, in the three relevant parameters, Δ*S*, Δ*w* and Δ*τ*_2_.

We assume that the two sub-networks store the same number *p* of memory patterns (with the same sparsity *a*), and that all the connections already encode these *p* patterns, as a result of a learning phase which is not modelled. We have seen in a previous study [29] that a differentiation Δ*S* has important dynamical implications during learning itself, but here we imagine learning to have already occurred. For a statistical study, we take the activity patterns to have been randomly generated with the same statistics, therefore any correlation between pattern *µ* and *ν* is random, and randomly different if calculated over each sub-network. These restrictive and implausible assumptions – they discard for example the possibility of structured associations between frontal and posterior patterns of different numerosity, statistics and internal non-random correlations – are needed to derive solid quantitative conclusions at the level of network operation, and might be relaxed later in more qualitative studies.

### 2.1 Connectivity in the differentiated network

For the statistical analysis, carried out through computer simulations, to be informative, the structure of the network model and in particular its connectivity have to be chosen appropriately. First, each sub-network should have the same number of units (half the total) and each unit the same number of inputs, for the comparisons between different conditions to be unbiased by trivial factors. Second, each sub-network should be allowed to determine, to some extent, its own recurrent dynamics, which requires the inputs onto each unit from the two halves not to be equal in strength, which would lead to washing away any difference, effectively, at each recurrent reverberation.

We then set the connection between units *i* and *j*, in their tensorial states *k* and *l*, as

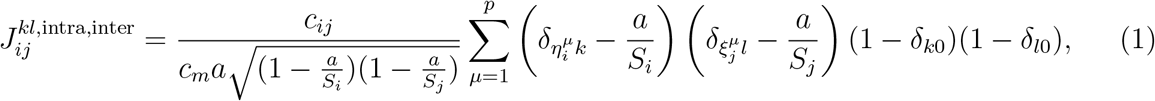

where {*c*_*ij*_} is a sparsity {0, 1} matrix that ensures that Potts unit receives *c*_*m*_ *intra* inputs from other units in the same sub-network and also receives *c*_*m*_ *inter* inputs from units of the other sub-network. Note that the number of Potts states of each unit, *S*, may depend on which sub-network the unit belongs to.

The partially differential dynamics is obtained by setting the strength coefficients as

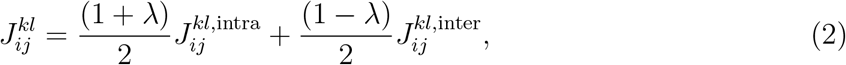

where the parameter *λ* ∈ [−1, 1] controls the relative strength of two terms. For *λ* = 0.0, the connectivity matrix becomes homogeneous and we cannot distinguish the two sub-networks from connectivity alone. If *λ* = 1.0, each sub-network is isolated from the other. For values of *λ* between 0 and 1, the recurrent connections within a sub-network prevail over those from the other sub-network, generating partially independent dynamics. We set *λ* = 0.5 as our reference value.

## 3 Results

We assume that the attractors of the frontal network have been associated one-to-one with those of the posterior network, via Hebbian plasticity, during a learning phase, which we do not model. When there is no external stimulus, e.g. when modelling creative thinking and future imaging, the network can sustain *latching* dynamics, i.e. it can hop from state to state, as in Fig. 1, provided its activity is appropriately regulated by suitable thresholds, as we have reported elsewhere [27]. Such spontaneous dynamics of the entire network might be led to a different extent by its frontal and posterior halves, depending on their characteristic parameters.

**Figure 1:**
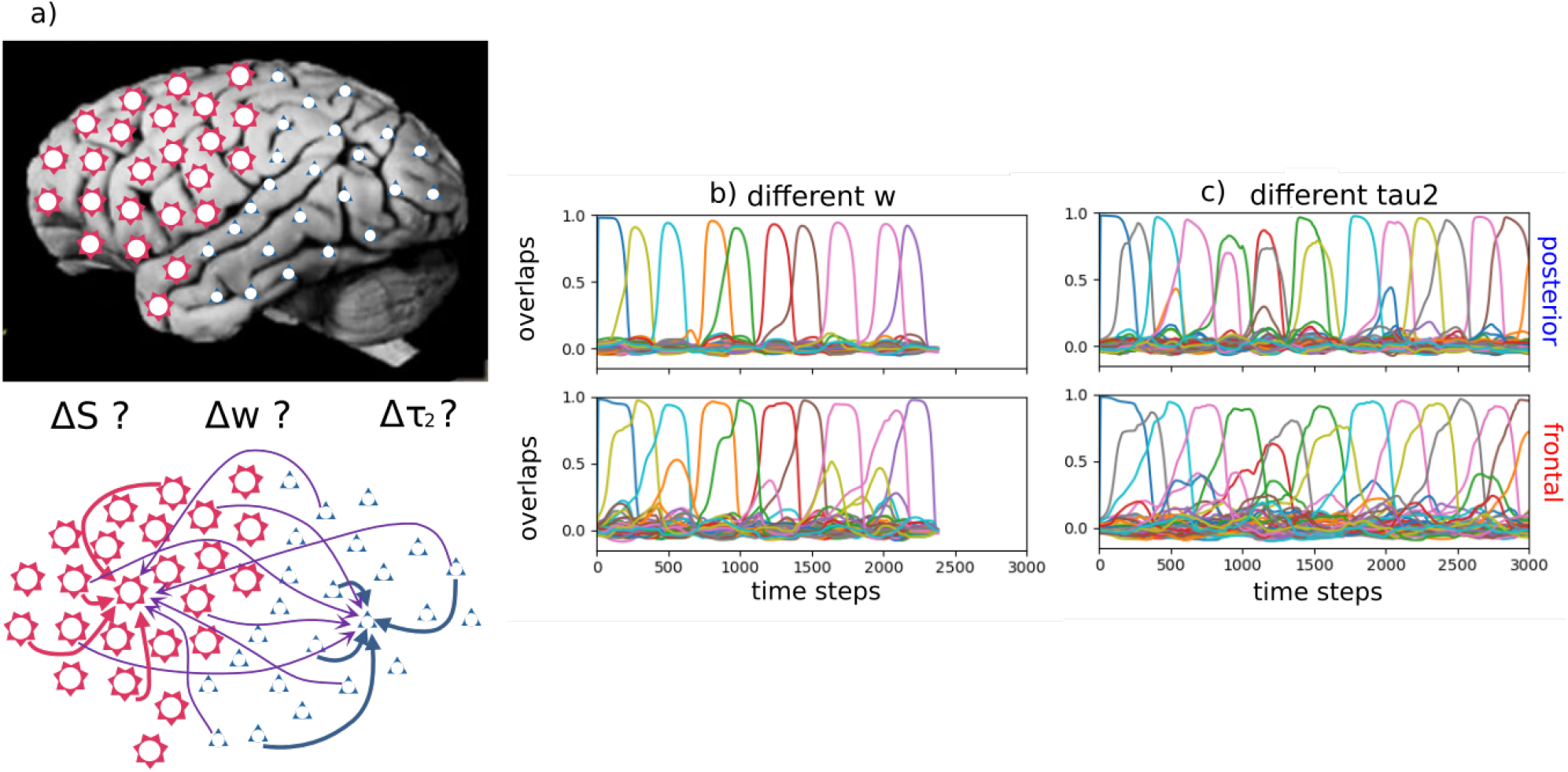
The differentiated network and examples of latching sequences. **(a)**: The differentiated network is comprised of frontal and posterior halves, in each of which units receive the same number of inputs from both halves, but not of the same average strength. **(b) and (c)**: The latching sequences are very similar if extracted from the posterior (upper panels) or the frontal sub-network (bottom panels). In (b), parameters are set as in Fig. 2e. In (c), parameters are set as in Fig. 3c.

In order to quantify the relative influence of the two sub-networks on the latching sequences produced by the hybrid Potts model, we look at whether the actual occurrence of each possible transition depends on the correlations, computed separately in the frontal and posterior parts, between the two patterns before and after the transition.

For the randomly correlated patterns used here, the correlations are relatively minor, but they can be anyway quantified by two quantities, *C*_*as*_ and *C*_*ad*_ ([30, 31]), that is, the fraction of active units in one pattern that are co-active in the other and in the same, *C*_*as*_, or in a different state, *C*_*ad*_. In terms of these quantities, two memory patterns are highly correlated if *C*_*as*_ is larger than average and *C*_*ad*_ is smaller than average, and we can take the difference *C*_*ad*_ − *C*_*as*_ as a simple compact indicator of the “distance” between the two patterns.

How strongly are transitions in a latching sequence driven by pattern correlations in each subnetwork? To measure this, we take the weighted average of *C*_*as*_ and *C*_*ad*_ with the weights given by latching sequences; that is, we compute

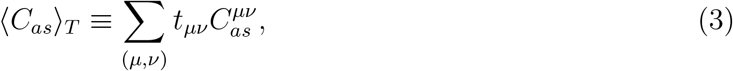

(and analogously for ⟨*C*_*ad*_⟩_*T*_) where the sum Σ_(*µ,ν*)_ runs over all possible pairs of memories and *t*_*µν*_ is the normalized frequency of latching transitions for the pair *µ, ν*: Σ _(*µ,ν*)_ *t*_*µν*_ = 1. This average is compared with the “baseline” average, e.g.,

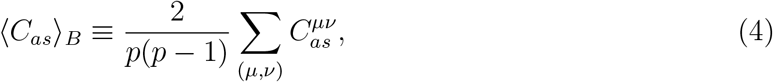

independent of the transitions, where *p* is the number of stored memories in the network. The comparison between the two averages, ⟨*C*_*as*(*d*)_⟩_*T*_ and ⟨*C*_*as*(*d*)_⟩_*B*_, is one index of how strongly latching sequences are related to correlations between patterns in one of the two sub-networks.

Second, based on the hypothesis that the frequency of transitions tends to decrease exponentially with the distance between the two patterns, as defined above, we look for the linear regression between the logarithm of the normalized transition frequency, log(*t*), and the distance *C*_*ad*_ − *C*_*as*_.

We first consider a case when all the macroscopic parameters are equal between the two subnetworks, while the connection parameter is set as *λ* = 0.5. In this case, the intra-connections (within each sub-network) are 3 times, on average, as strong as the inter-connections (between the two sub-networks), but the two halves are fully equivalent, or Not Differentiated (ND). With the appropriate parameters, in particular the feedback *w*, we find that the network as a whole shows robust latching and that latching sequences in each sub-network are well synchronized with each other: the two sub-networks essentially latch as one. Comparing latching dynamics in two sub-networks, we find that latching is largely driven by correlations between patterns, in either half or in both, as found previously [30]. This can be seen, leftmost bars of Fig. 2a and Fig. 2b, by the higher value of ⟨*C*_*as*_⟩_*T*_ relative to ⟨*C*_*as*_⟩_*B*_, and vice versa for *C*_*ad*_, in the ND case. Correlations in the two sub-networks appear to contribute equally to determine latching sequences, as expected. This is confirmed by the similar negative slopes in the two scatterplots of Fig. 2c.

**Figure 2:**
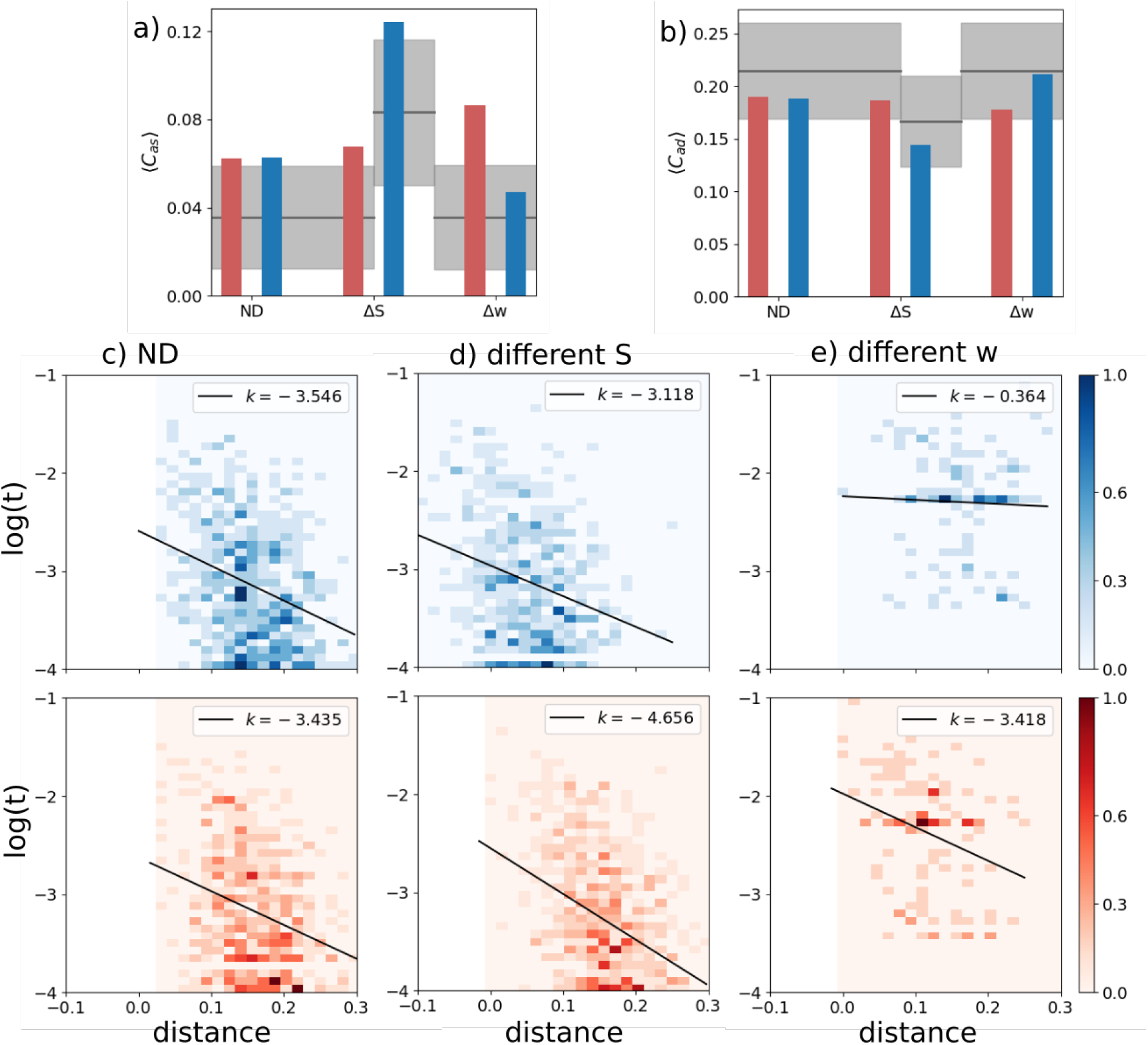
A latching frontal network leads a non-latching posterior network. Red indicates the frontal and blue the posterior network in this and other figures. **(a) and (b)** The transition-weighted averages of *C*_*as*_ and *C*_*ad*_ are compared to their baseline values for three cases: no difference between the two networks (ND, leftmost bars), a difference in *S* (Δ*S*, middle bars) and a difference in *w* (Δ*w*, rightmost bars). The gray horizontal line and shaded area indicate the baseline average and its standard deviation. **(c), (d) and (e)** Scatterplots of (log) transition frequencies between individual patterns pairs versus their distance, for the three conditions. The darkness of color indicates the number of pairs at each combination of abscissa and ordinate. For the ND condition, parameters are set as *w*_*p*_ = *w*_*f*_ = 1.1, *S*_*p*_ = *S*_*f*_ = 7. For the other conditions, the parameters of the frontal network are kept the same as in the ND condition, while the parameters of the posterior sub-network are set as *S*_*p*_ = 3 and *w*_*p*_ = 0.6, respectively, in (d) and (e).

**Different** *S*. We now examine a case in which the two networks share the same values of all but one parameter: the number of Potts states, *S*. When the posterior network has fewer states (*S* = 3 instead of the reference value, 7), the baselines for both *C*_*as*_ and *C*_*ad*_ are shifted, above and below, respectively, but their transition-weighted values are similarly positioned, above and below the respective baselines, as in the frontal network. Also in terms of the second indicator, the scatterplot of Fig. 2d shows rather similar slopes, with only a modest quantitative “advantage” for the frontal network (in red), which can be said to lead the latching sequence somewhat more than the posterior one. One should note that, with these parameters, both sub-networks would latch if isolated.

**Different** *w*. In contrast to the two cases above, ND and Δ*S*, we see a major difference between the two sub-networks if it is the *w* parameter which is lower for the posterior network (the rightmost bars of Figs. 2a,b). In this case, it is obviously the correlation structure of the frontal patterns, not of the posterior ones, that dominates in determining latching sequences. This is also evident from the very different slopes, *k*, in the scatterplot of Fig. 2e. With the lower value *w* = 0.6 chosen for the posterior sub-network, this time it would not latch, if isolated. Note that to preserve its latching, and for it to be a clear single sequence, we would have to set *w* at almost the same value as for the frontal sub-network, unlike the case with the *S* parameter.

**And/or different** *τ*_2_. We now allow the adaptation timescale, *τ*_2_, to differ between two sub-networks. We first note that latching sequences between the two networks are remarkably well synchronized despite their different adaptation timescales (Fig. 1c). If isolated, the two sub-networks would each latch at a pace set by its own *τ*_2_. Their synchronization thus shows that, even with this relativity weaker connectivity coupling (inter-connections 1/3 of the average strength of the intra-connections) the two halves are willing to compromise, and latch at some intermediate pace, close to the one they sustained when *τ*_2_ was not differentiated.

Furthermore, latching sequences are affected predominantly by frontal correlations rather than posterior ones. In Fig. 3, we show two cases: the two sub-networks have two different adaptation timescales; and in the second case also different *w*. We see a moderate effect if *τ*_2_ is the only parameter that differs between the two. Note that in this case the posterior sub-network, if isolated, would latch.

**Figure 3:**
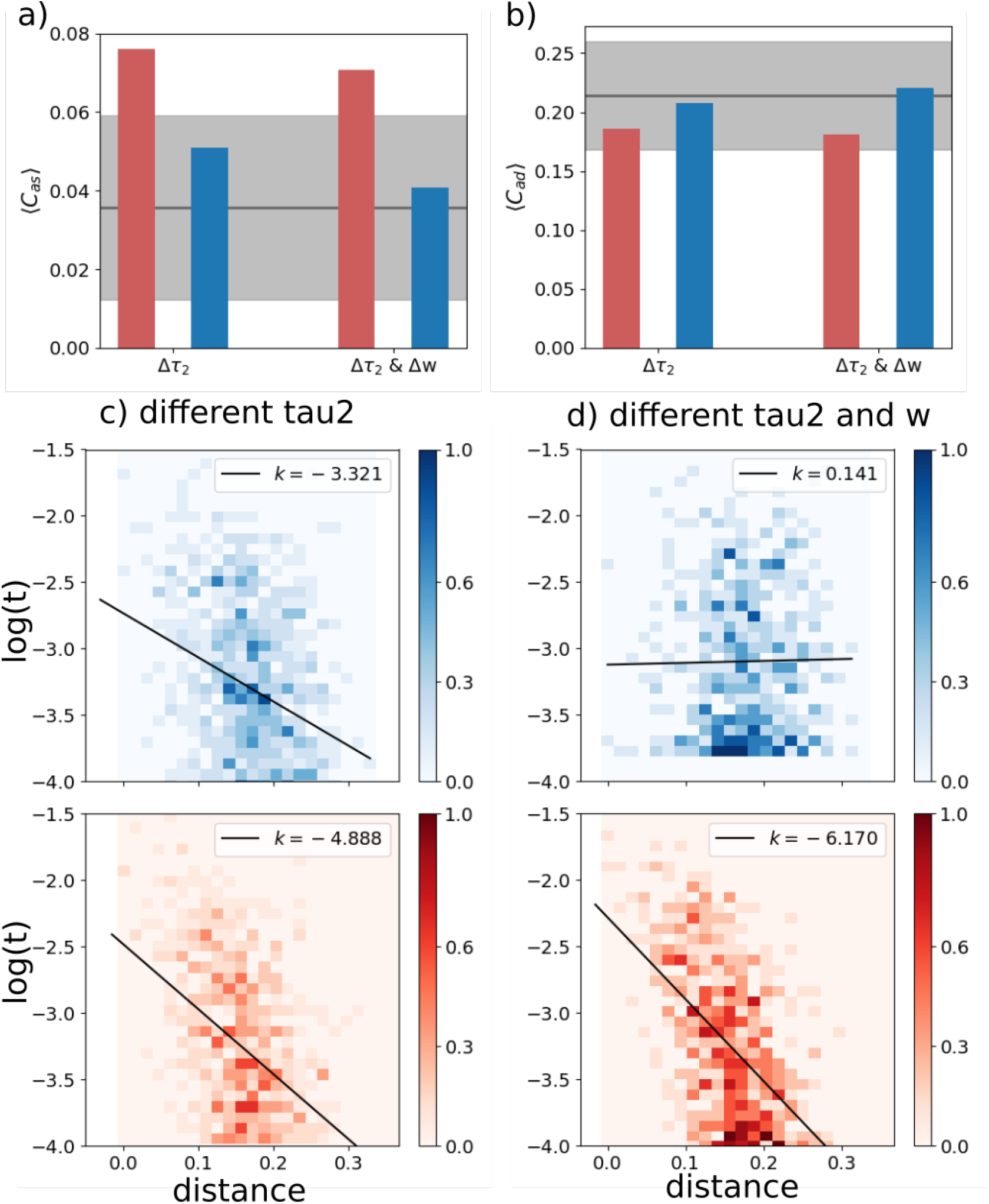
The frontal sub-network is even more dominant with slower adaptation. Color code and meaning are the same as in Fig. 2. **(a) and (b)** Transition-weighted averages of *C*_*as*_ and *C*_*ad*_ versus their baselines are shown for two conditions: only *τ*_2_ is different and both *w* and *τ*_2_ are different. In both conditions, *τ*_2_ is 100 for the posterior network and 400 for the frontal network. In the Δ*w* condition, *w* is 0.6 for the posterior network and 1.1 for the frontal network. **(c) and (d)** Log-transformed transition frequencies between individual patterns pairs versus their distance.

The effect is most pronounced if *w* is also lowered to *w* = 0.6 for the posterior sub-network, as is evident from the weak positive slope *k* it shows, see Fig. 3d. In this case it would not latch if isolated.

We have also inverted the *τ*_2_ difference, making the posterior sub-network, still with a lower *w*, slower in terms of firing rate adaptation. In this case (not shown) latching is virtually abolished, showing that the parameter manipulations do not simply add up linearly.

### 3.1 Lesioning the network

To model lesions in either sub-network, we define a procedure that still allows us to compare quantities based on the same number of inputs per unit, etc. The procedure acts only on the relative weights of the connections (through *λ*), which are modulated while keeping their average for each receiving unit always to 1*/*2. Other parameters of the network are set in such a way that the frontal sub-network leads the latching sequences and that lesions do not push the network into a no-latching phase: the self-reinforcement parameter is set as *w* = 0.7 for the posterior sub-network and *w* = 1.2 for the frontal one, while *S* and *τ*_2_ are set as specified in Table 1 and thus take the same value for both sub-networks. For “healthy” networks, we use *λ* = 0.5 in Eq. (2), meaning the intra-connections (within the frontal and within the posterior half) are 3 times, on average, as strong as the inter-connections (between frontal and posterior halves). For lesioned networks, we use smaller values of *λ* than 0.5 for their input connections: the smaller the value is, the stronger the lesion is. So, for example, a frontal lesion with *λ* = 0.2 implies that its recurrent weights are weighted by a factor 0.6 (instead of 0.75) and the weights from the posterior sub-network by a factor 0.4 (rather than 0.25), i.e. the internal weights are only 1.5 times those of the interconnections. The posterior sub-network in this case has the same weights as the control case.

**Table 1:** Parameters of the network.

| Symbol | Meaning | Default value |
| --- | --- | --- |
| $N$ | number of Potts units | 256 |
| $c_m$ | number of presynaptic units | 50 |
| $S$ | number of states per unit | 7 |
| $p$ | number of memory patterns | 50 |
| $a$ | sparsity of patterns | 0.25 |
| $\lambda$ | relative coupling strength | 0.5 |
| $U$ | global threshold | 0.1 |
| $\tau_2$ | adaptation timescale | 200 |
| $w$ | self-reinforcement term | 1.1 |
| $\beta$ | inverse “temperature” | 11 |

We then quantify the effect of the lesions with the slopes in the scatterplots as before, but also with an entropy measure. The entropy at position *z* in a latching sequence measures the variability of transitions encountered at that position, across all sequences with the same starting point. It is computed as

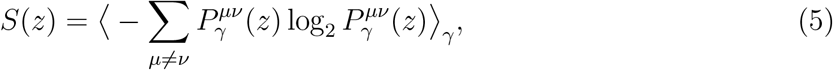

where 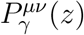 is the joint probability of having two patterns *µ* and *ν* at two consecutive positions *z* and *z* + 1 relative to the cued pattern *γ* in a latching sequence, and ⟨·⟩_*γ*_ means that we average the entropy across all the *p* patterns that are used as a cue. Note that if all transitions were incurred equally, asymptotically for large *z*, the entropy would reach its maximum value *S*_*∞*_ = log_2_[*p*(*p* − 1)] (with *p* patterns stored in memory and available for lathing). Therefore exp{[*S*(*z*) − *S*_*∞*_] ln(2)} is an effective measure of the fraction of all possible transitions that the network has explored at position *z*, on average.

In terms of the slopes in the scatterplots, we see that posterior lesions do not have a major effect, while frontal lesions reduce the relation between the probability of individual transitions and the correlation between the two patterns, particularly in the frontal sub-network where it was strong in the “healthy” case (Fig. 4).

**Figure 4:**
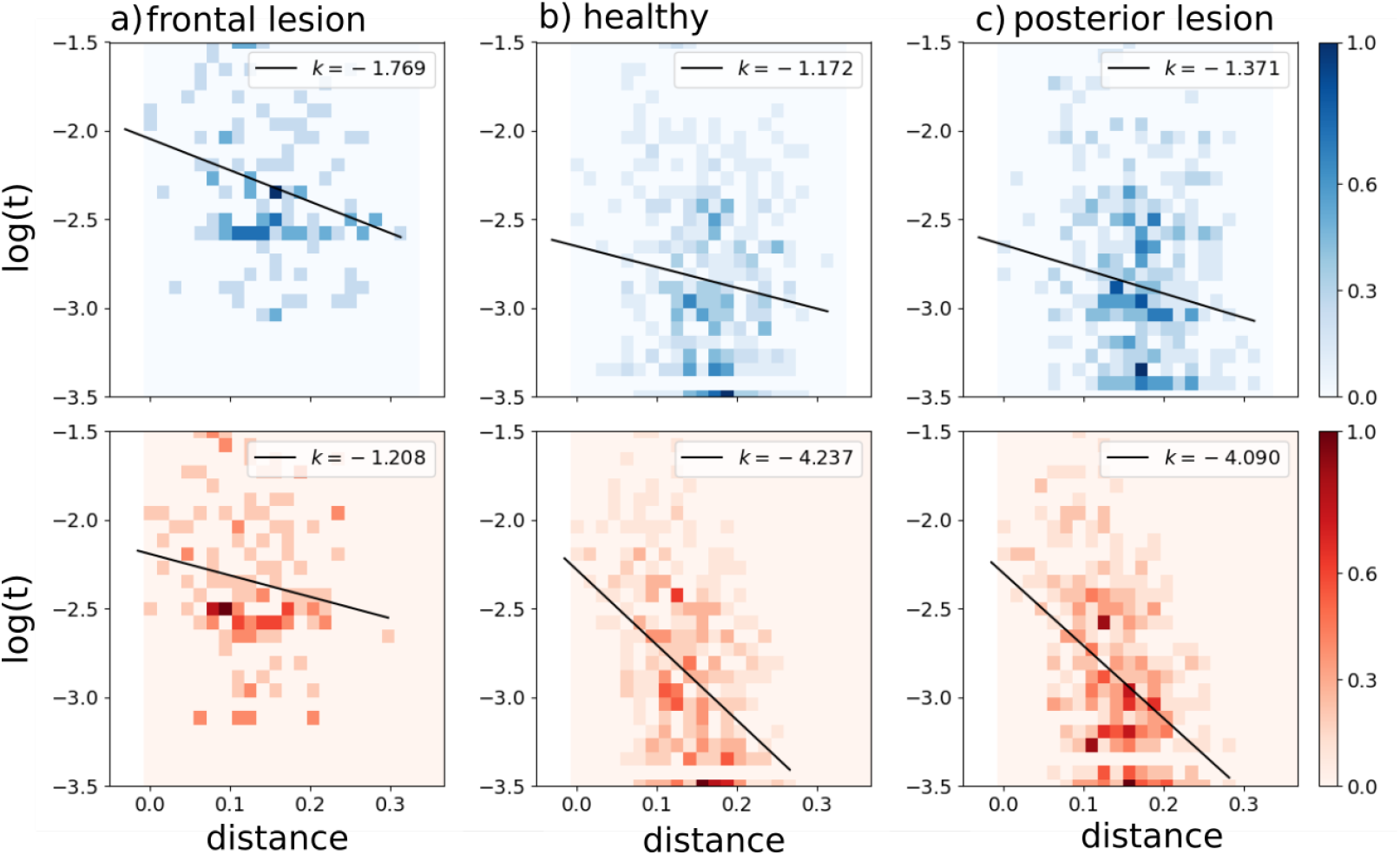
Correlations between transition frequency and pattern distance are shown for a network with frontal lesions (a), for a healthy network (b) and for a network with posterior lesions (c). Lesions are modelled by setting *λ* = 0.2 (see main text). The self-reinforcement parameter is set as *w* = 1.2 for the frontal sub-network and *w* = 0.7 for the posterior one.

In terms of entropy, we see that lesions in the posterior sub-network do not affect the entropy curve, relative to that for the healthy network (Fig. 5). Lesions in the frontal subnetwork, however, tend to restrict the sequences to a limited set of transitions, leading to a marked reduction in the fraction of possibilities explored by the lesioned network.

**Figure 5:**
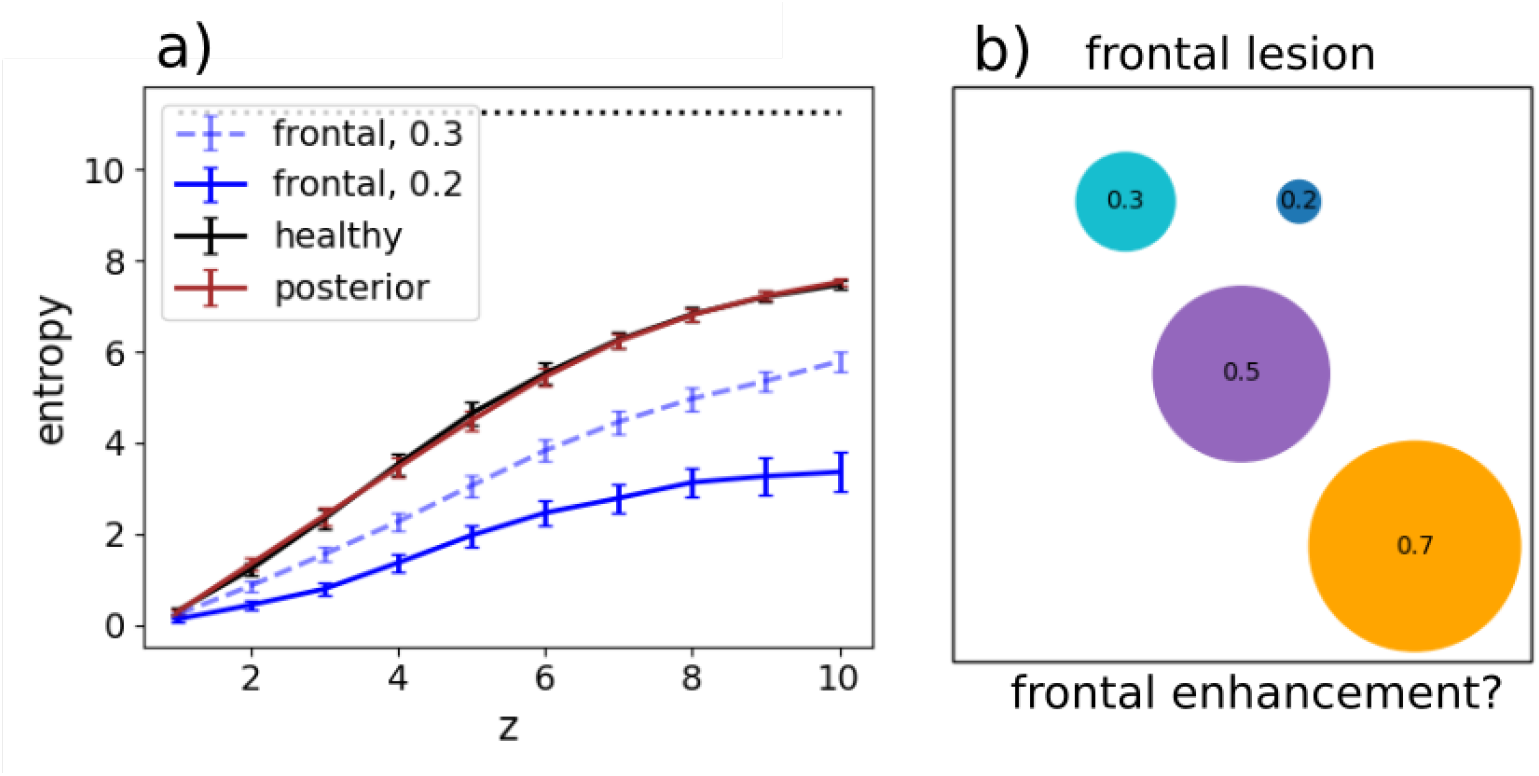
**(a)** The entropy *S*(*z*) and its standard error of the mean are shown for healthy (black), frontal-lesioned (blue) and posterior-lesioned (red) networks. Lesions are implemented by setting *λ* = 0.2 for solid curves, whereas the dashed blue curve is for a milder lesion in the frontal network (*λ* = 0.3). The black horizontal line indicates the asymptotic entropy value for a completely random sequence generated from a set of *p* = 50 patterns. The self-reinforcement parameter is set as *w* = 1.2 for the frontal network and *w* = 0.7 for the posterior network. **(b)** A schematic view of the diversity of transitions expressed by latching sequences. Circles are centered around an arbitrary position, while their areas extend over a fraction 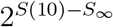 of the area of the square (which would correspond to an even exploration of all possible transitions, asymptotically). The large orange circle is obtained by setting *λ* = 0.7, thus modelling a sort of cognitive *frontal enhancement*, perhaps obtained with psychoactive substances.

Simulated frontal lesions, therefore, produce in our model two effects that, while not opposite, are not fully congruent either. The first, manifested in the reduced slope of Fig. 4a, is suggestive of a loss of coherence in individual transitions between brain states; the second, seen in the limited entropy of Fig. 5, indicates a restriction in the space spanned by the trajectories of spontaneous thought. To reconcile the two outcomes, we have to conclude that while less dependent on the similarity between the two patterns, or states, individual transitions are not really random, and some become in the lesioned network much more frequent than others, gradually veering from creative towards obsessive (or perseverative) thought.

## 4 Discussion

Simulating our model provides some insight about the conditions that may enable frontal cortex to determine the sequence of states in spontaneous thought dynamics. It is important, in assessing the computational findings, to distinguish what has gone into defining the model from what the model gives out in return. For example, much cognitive neuroscience research has been devoted to understanding the process of segmenting our ongoing experience into separate sub-events, or event segmentation [32]. Baldassano and colleagues [33] have recently demonstrated how brain activity within sub-events resembles temporarily stable activity patterns, dubbed “neural states” [34], which may be identified with those long posited to occur in the cortex of primates [35] and other species [36], from analyses of single-unit activity. This notion is conceptually similar to the Potts states in a latching sequence, but finding evidence that a continuous input flow is segmented into discrete or quasi-discrete states in the brain is a major achievement, whereas in the Potts network it is a straightforward outcome of the ingredients used to define the model in the first place. Interestingly, these neural states were found to occur on different timescales across regions, with more but short-lasting transitions in low-level (posterior) sensory cortices and fewer but longer-lasting transitions in higher-level (frontal/parietal) regions. Strikingly, for some of the higher order brain regions, neural state transitions appeared to overlap with behavioural measures of event boundary perception [37]. In our study, the central question is which portion of the differentiated model network controls the sequence of discrete event states. We have seen that three types of differentiation, each capturing some aspect of caudo-rostral cortical variation, bias sequence control towards the “frontal” half of the network, albeit with different effectiveness. A comparison across the three types of differentiation is inherently ill-defined, because Δ*S*, Δ*w* and Δ*τ*_2_ are all measured on different scales, but it is apparent that the first type has a much milder effect than the second, and the third is somewhere in between. The major effect seen with Δ*w* is likely due to the posterior network being unable to latch on its own, with the lower *w* value we have used. The lower *S* and *τ*_2_ values do not have much of an effect on latching *per se*. The three types of differentiation are of course not mutually exclusive, and it is plausible that in the real brain, if the model makes sense, their effect would be cumulative. They do not appear to add up linearly, though: we have mentioned that inverting the *τ*_2_ difference with respect to the *w* difference (i.e., making firing rate adaptation faster in the frontal sub-network) tends to abolish latching altogether, rather than reduce the frontal advantage in leading it.

A limitation of our study is that to compare the sub-networks on an even footing we have considered an artificial scenario in which activity patterns are only randomly correlated, and also there are *p* in each half network and they have been paired one-to-one during learning. Obviously in this scenario there is no benefit whatsoever if the network follows a frontally-rather than a posteriorly-generated sequence: they are equivalent, and both devoid of content. It will be therefore important, in future work, to understand whether the insights derived under these assumptions are applicable also to more plausible conditions, in which the frontal and posterior patterns are not paired one-to-one, and can take distinct roles, for example along the lines of the classic operator/filler (also denoted as role/filler) distinction [38]. In this more complex scenario, the frontal patterns, if they have to serve as operators, would “take” or be paired in certain cases to a single filler and in others to multiple fillers (and possibly to other operators, in a hierarchical scheme); but even if just to one, it would be one among several options, so the pairing scheme in long-term-memory would be considerably more complex than the one considered here.

A relevant cognitive construct we mention, only partially overlapping with that of operator, is that of a temporally-oriented *schema*. A schema is a regularity extracted from multiple experience, in which B follows A and is then followed by C, although the particular instantiation of A, B and C will be different every time [39]. Note that to be implemented in our network, the skeleton of the ABC representation would have to stay activated while the specific filling items A, B and C are specified, in succession, in the posterior cortex. Alternatively, ABC could be conceptualized as a short tight latching sequence. Clearly, more attention has to be paid to the possibility of formalizing these constructs in a future well-defined network model.

### Mind wandering and creativity

Within its present limitations, still our approach may offer insights relevant to the dynamics of state transitions in spontaneous cognition, such as those underlying mind wandering. Mind wandering occurs when attention drifts away from ongoing activities and towards our inner world, focusing for example on memories, thoughts, plans, which typically follow one another in a rapid, unconstrained fashion [40, 41]. The dynamics governing the flow of thoughts can indeed be described as latching (see also [6]).

Mind wandering is known to engage the Default Mode Network (DMN), a set of interconnected brain regions, spanning from posterior, temporal, and frontal cortices [42, 43, 44, 45, 41, 46], underlying introspection and spontaneous (endogenously triggered) cognition. Ciaramelli and Treves [6] and McCormick et al. [47] have proposed that the prefrontal cortex, especially in its ventral-medial sectors (vmPFC) might support the initiation (internal triggering) of mind-wandering events. Indeed, recent MEG findings show that activity in the vmPFC precedes (presumably drives) hippocampal activity during (voluntary) scene construction and autobiographical memory retrieval ([48]; see also [49, 50]), and this region may play a similar role during spontaneous cognition. Indeed, damage [28, 51] or inhibition [52, 53] of the vmPFC (but not the hippocampus; [47]) reduce the frequency of mind-wandering.

On one view, vmPFC initiates event construction by activating schemata (about the self, or common events) that help collect relevant details that the hippocampus then binds in coherent, envisioned scenes ([54]; see also [55, 56]). Consistent with the schema hypothesis, vmPFC (but not hippocampal) patients are particularly impaired in event construction when the task benefits from the activation of the self schema ([57]; [58]), and are not impaired when the need for self-initiation is minimized [59]. vmPFC may also govern schema-congruent transitions between successive scenes of constructed events based on event schemata (scripts) ([46]; [60]), which may explain why vmPFC patients are particularly poor at simulating extended events as opposed to single moments selected from events [28, 61]. The results from our computational simulations accord with and complement this view. Lesioning the frontal (but not the posterior) sector of the network led to more random state transitions, less dependent on the correlation between patterns, and also led to shorter-lasting sequences, that fade out after fewer state transitions. This pattern of findings is expected if transitions in thought states were not guided by schematic knowledge, making them less coherent in content and self-exhausting.

A second effect we observed is a reduced entropy following lesions in the frontal (but not posterior) half of the network, which indicates that the trajectories of state transitions were confined in a limited space, as if mind wandering lost its ‘wandering’ nature to become more constrained, with recurring thoughts characteristic of the perseverative responses long observed in prefrontal patients; suggesting that vmPFC patients, in addition to an impaired activation of relevant schemata, also fail in flexibly *de*activating current but no longer relevant ones [39].

The most characteristic memory deficit following vmPFC damage is confabulation, the spontaneous production of false memories. Confabulations often involve an inability to inhibit previously reinforced memory traces [62]. For example, confabulators can falsely endorse personal events as true because these were true in the past (e.g., that they just played football while in fact they used to play football during childhood). If presented with modified versions of famous fairy tales to study, confabulators tend to revert to the original versions of the stories in a later recall phase [63]. Similarly, during navigation, confabulators may get lost because they head to locations they have attended frequently in the past, instead of the currently specified goal destination [64].

The inability to flexibly switch between relevant time schemata and memory traces has been linked to reduced future thinking and reduced generation of novel scenarios in prefrontal patients ([65]; see also [28]), who admitted they found themselves bound to recast past memories while trying to imagine future events. More in general, prefrontal lesions impair creativity. There is interaction between the DMN and the fronto-parietal control network while generating (DMN) and revising (fronto-parietal network) creative ideas ([66, 67]). Bendetowicz et al. found that damage to the right medial prefrontal regions of the DMN affected the ability to generate remote ideas, whereas damage to left rostrolateral prefrontal region of the fronto-parietal control network spared the ability to generate remote ideas but impaired the ability to appropriately combine them.

Note, however, that the originality associated with creative ideas can be conceived as disrupting the automatic progression from a thought to the one most correlated to it. Fan et al. ([68]) had participants perform a creative writing task, and indeed found the *semantic distance* between adjacent sentences to be positively correlated with the story originality. Also, semantic distance was predicted by connectivity features of the salience network (e.g., the insula and anterior cingulate cortex) and the DMN. Green et al. ([69]) have also reported a putative role of mPFC (BA 9/10) in connecting semantically distant concepts during abstract relational integration. In a following study ([70]), mPFC activity was found to vary monotonically with increasing semantic distance between abstract concepts, even when controlling for task difficulty. Indeed, preliminary evidence from patients with vmPFC lesions is indicative of a greater global semantic coherence in speech compared to healthy participants (Stendardi et al., in preparation). These results align with our finding that a lesion of the frontal component of the network produces a reduction in entropy, making latching dynamics “less creative”; but not, *prima facie*, with the reduced slope in Fig.4a, which indicates that the lesion would produce more random transitions, frequent also among distant patterns. The apparent contradiction can be reconciled by noting that, as seen above, *individual* random transitions can still result in reduced entropy, if they tend to recur perseveratively within a sequence; and also that *semantic* coherence may reflect pattern correlation in posterior rather than frontal cortices, whereas it is logical/syntactic consequentiality that is expected to be impaired by random frontal transitions. In fact, in our model lesion, the decreased slope in the frontal sub-network seen in Fig.4a (more random transitions) is accompanied by a slightly increased slope, suggestive of more semantic coherence, posteriorly.

Clearly, a major refinement of our approach is required, before these suggestions can be taken seriously, and articulated in a more nuanced view of how operating along the time dimension may be coordinated across cortical areas.

## Acknowledgments

This work was supported by PRIN grant 20174TPEFJ “TRIPS” to EC and AT. We are grateful for early discussions with Massimiliano Trippa.

## Data availability

The simulation code is available in the OSF repository at the link (to be made public upon acceptance) https://osf.io/fk5uz/?view_only=55fd1a67077846c8b44dd6e1d3933831.

## Conflict of interest

The authors declare no conflict of interest.

## Author contributions

KIR and AT conceived and developed the computational project as a means of articulating the neuropsychological perspective put forward by EC and DS. KIR run most of the simulations, which were complemented by others developed by AB. DS spurred the analysis of coherence. KIR drafted this report and all coauthors contributed to refine and complete it.

## A Potts model details

A Potts neural network is an autoassociative memory network comprised of *N* Potts units, which model patches of cortex as they contribute to retrieve distributed long-term memory traces addressed by their contents [27]. Each Potts unit has *S* active states, indexed as 1, 2, · · ·, *S*, representing local attractors in that patch, and one quiet state, the 0 state. The *N* units interact with each other via tensor connections, that represent associative long-range interactions through axons that travel through the white matter [7], while local, within-gray-matter interactions are assumed to be governed by attractor dynamics in each patch. The values of the tensor components are pre-determined by the Hebbian learning rule, which can be construed as derived from Hebbian plasticity at the synaptic level [25]

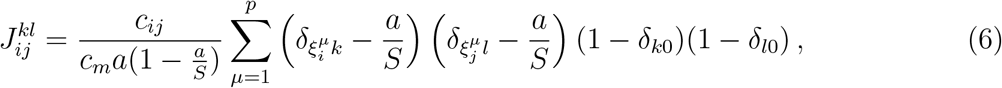

where *c*_*ij*_ is either 1 if unit *j* gives input to unit *i* or 0 otherwise, allowing for asymmetric connections between units, and the *δ*’s are the Kronecker symbols. The number of input connections per unit is *c*_*m*_. The *p* distributed activity patterns which represent memory items are assigned, in the simplest model, as composition of local attractor states 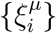 (*i* = 1, 2, · · ·, *N* and *µ* = 1, 2, · · ·, *p*). The variable 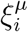 indicates the state of unit *i* in pattern *µ* and is randomly sampled, independently on the unit index *i* and the pattern index *µ*, from {0, 1, 2, · · ·, *S*} with probability

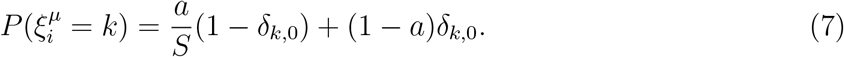

Constructed in this way, patterns are randomly correlated with each other. We use these randomly correlated memory patterns 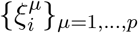 in this study. The parameter *a* is the sparsity of patterns – fraction of active units in each pattern; the average number of active units in any pattern *µ* is therefore given by *Na*.

Local network dynamics within a patch are taken to be driven by the “current” that the unit *i* in state *k* receives

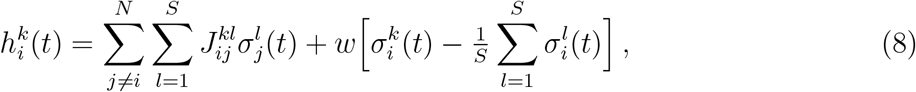

where the local feedback *w*, introduced in [30], models the depth of attractors in a patch, as shown in [25] – it helps the corresponding Potts unit converge to its most active state. The activation along each state for a given Potts unit is updated with a *soft max* rule

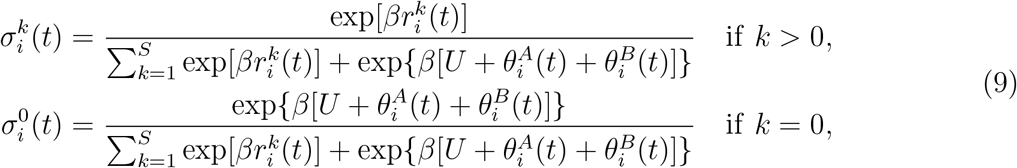

where *U* is a fixed threshold common for all units and *β* is an effective inverse “temperature” (noise level). Note that 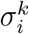 takes continuous values in (0, 1) and that 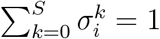 for any *i*. The variables 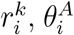 and 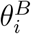 parametrize, respectively, the state-specific potential, fast inhibition and slow inhibition in patch *i*. The state-specific potential 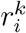 integrates the state-specific current 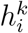 by

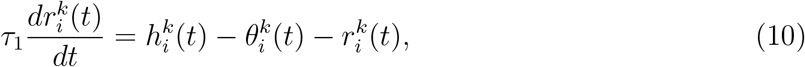

where the variable 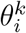 is a specific threshold for unit *i* and for state *k*.

Taking the threshold 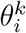 to vary in time to model adaptation, i.e. synaptic or neural fatigue selectively affecting the neurons active in state *k*, and not all neurons subsumed by Potts unit *i*

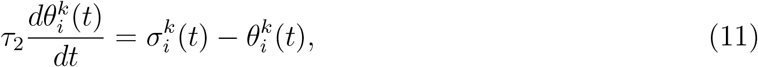

the Potts network additionally expresses latching dynamics, the key to its possible role in modelling temporal schemata.

The unit-specific thresholds 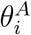 and 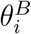 describe local inhibition, which in the cortex is relayed by at least 3 main classes of inhibitory interneurons [71] acting on GABA_A_ and GABA_B_ receptors, with widely different time courses, from very short to very long. Formally in our model, 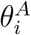 denotes fast, GABA_*A*_ inhibition and 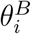 denotes slow, GABA_*B*_ inhibition and they vary in time in the following way:

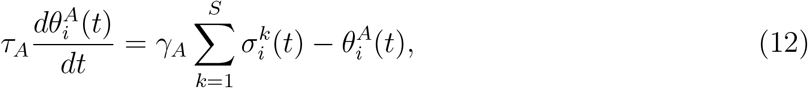

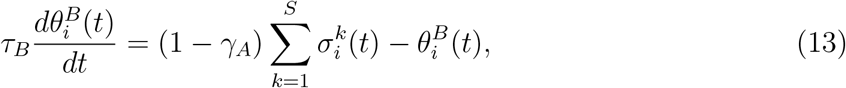

where one sets *τ*_*A*_ *< τ*_1_ ≪ *τ*_2_ ≪ *τ*_*B*_ and the parameter *γ*_*A*_ sets the balance of fast and slow inhibition. Specifically in this work, we set these parameters as *τ*_*A*_ = 10, *τ*_*B*_ = 10^5^, *τ*_1_ = 20 and *γ*_*A*_ = 0.5.

## B Simulation details

We have used an asynchronous updating, where one unit is updated at a time with a random order. Updating all Potts units in the network once is our measuring unit of simulation time: all timescales of the model are measured with this unit. We stop the simulation after updating the entire network 10000 times (except for Fig. 5, see next paragraph). Then, we cut out the first 3 patterns in the sequence to remove the effect of initialization. Every stored memory is used as a cue with its full representation.

In order to compute the probability 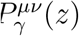 in Eq. (5), we have run *p* × 1000 simulations for each condition. For each memory pattern, we take 40% of its active units and flip them into different states. We prepare 1000 corrupted versions of each memory by repeating this procedure 1000 times. Each of these corrupted versions is used as a cue in each simulation, which is terminated after 12 transitions.

Unless specified explicitly, parameters of the Potts model are set as in Table 1.

## Notes

### Competing Interest Statement

The authors have declared no competing interest.

